## Supplementary figures and images for "Microbial colonization induces histone acetylation critical for inherited gut-germline-neural signaling"

### Supplemental Figures 1-7

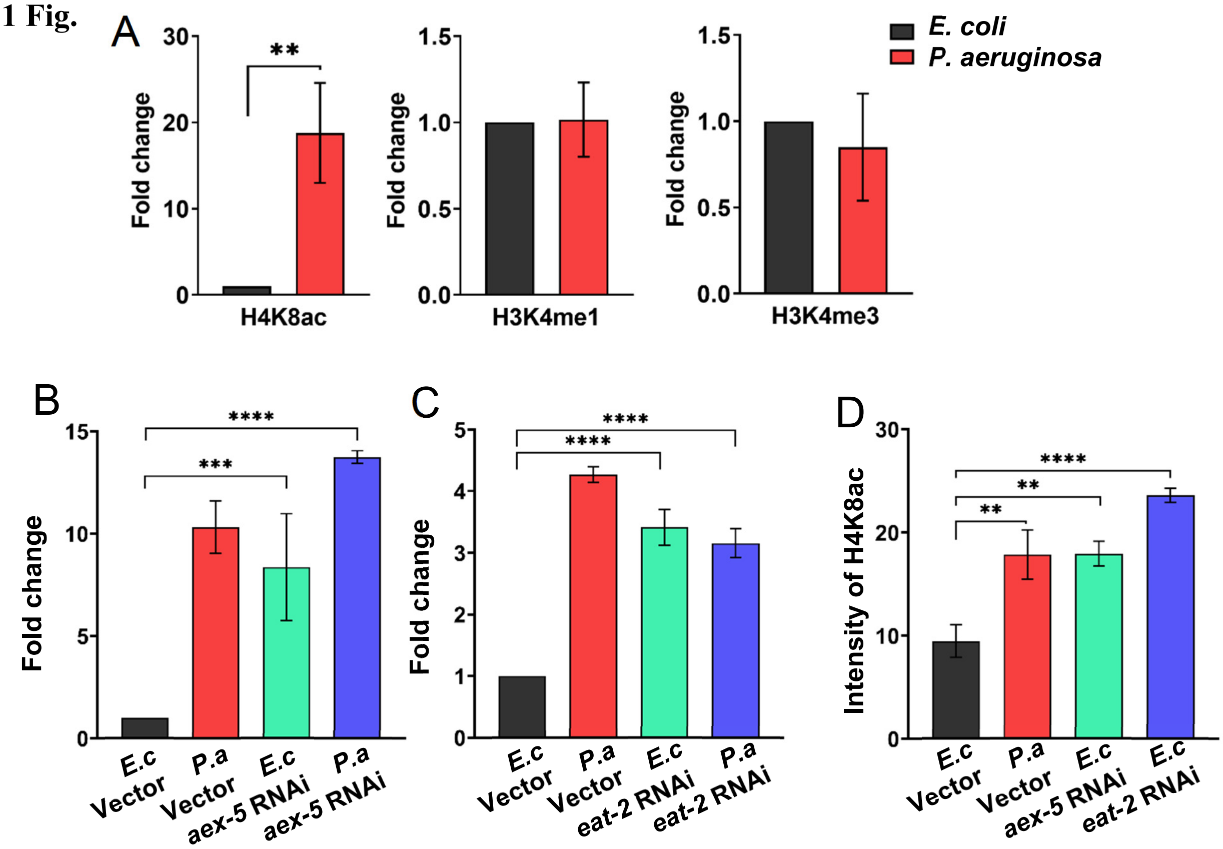

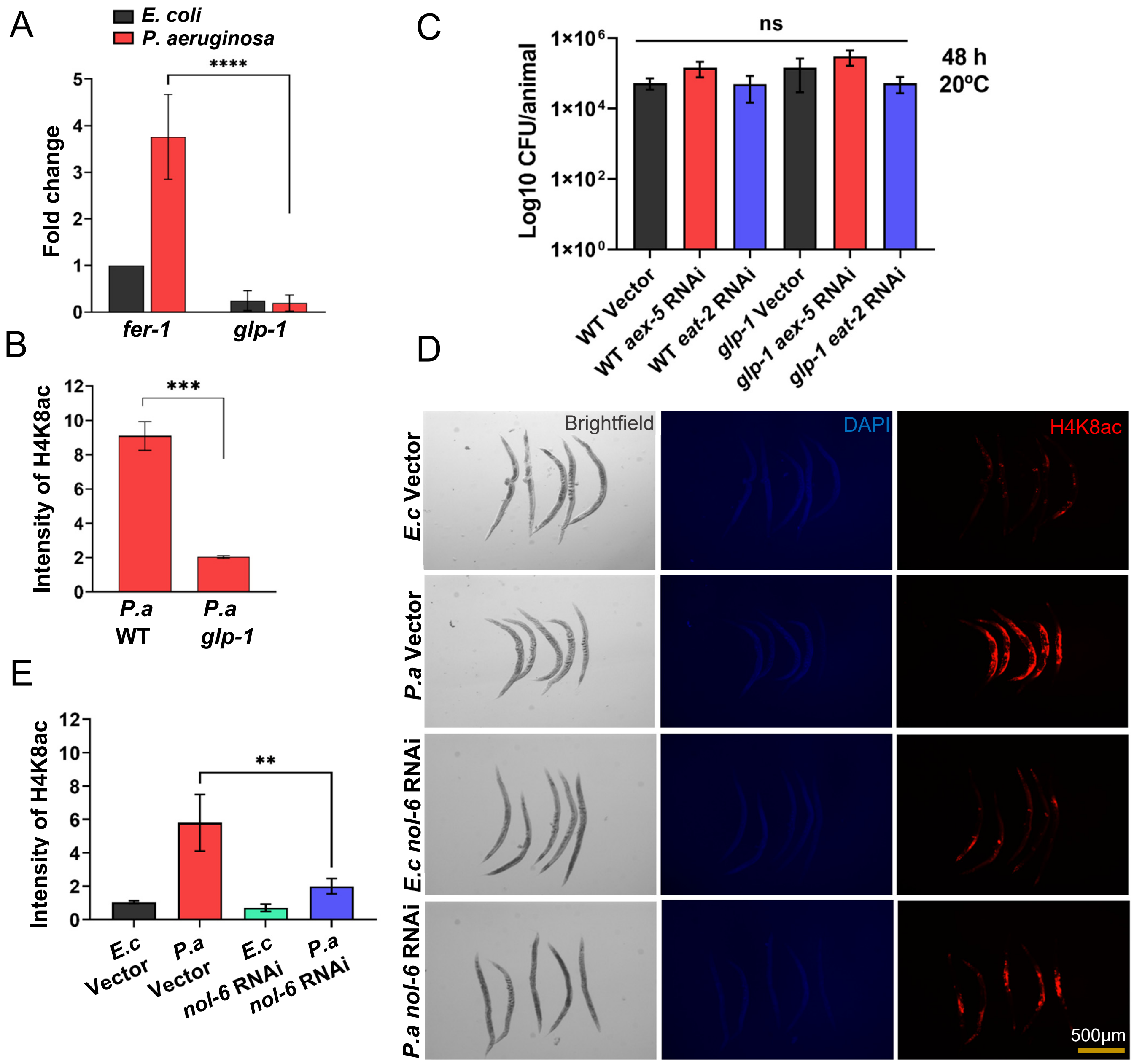

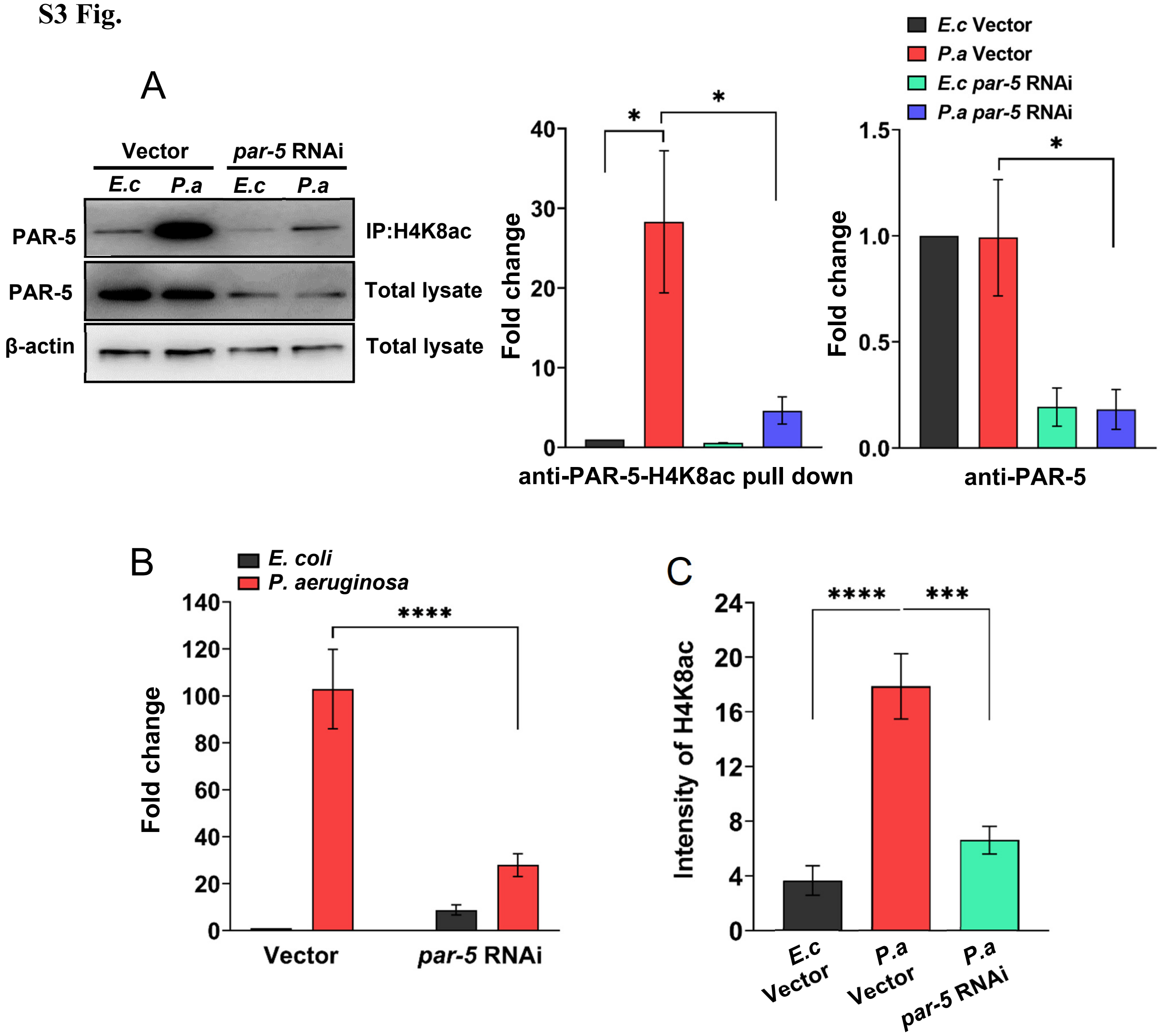

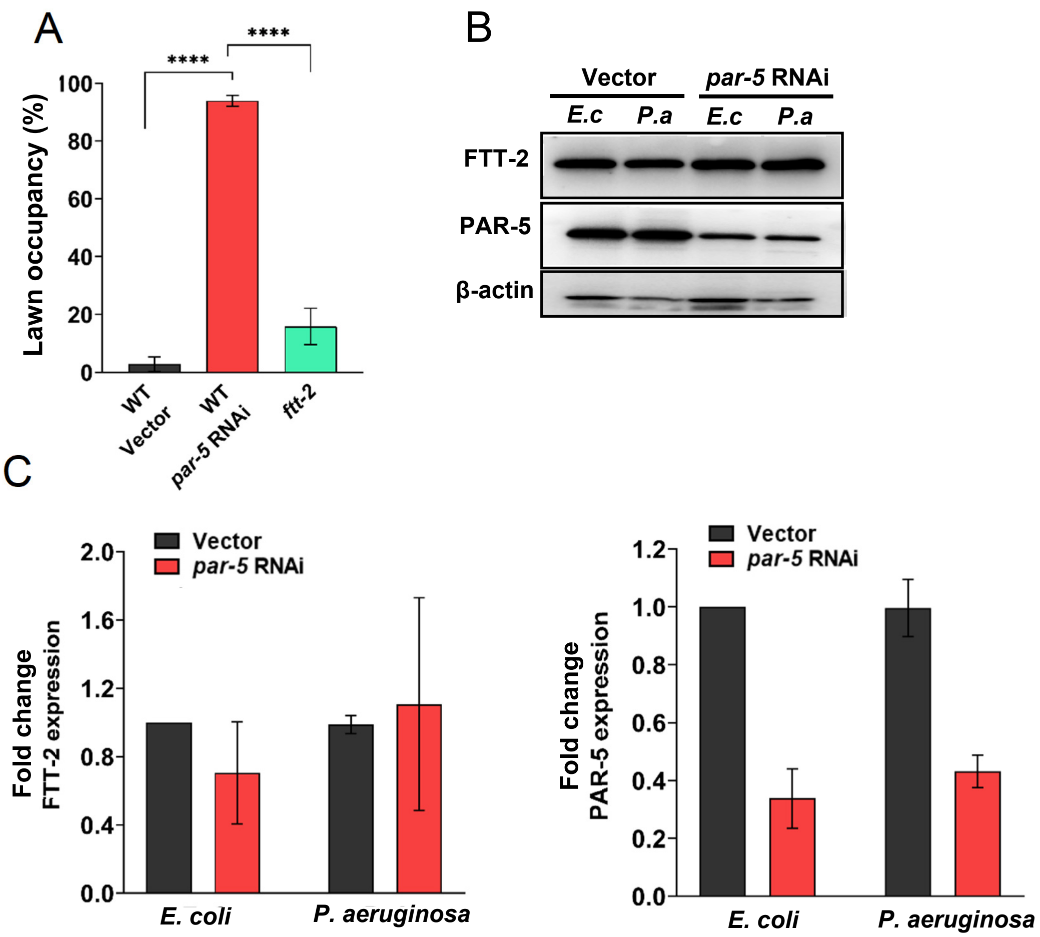

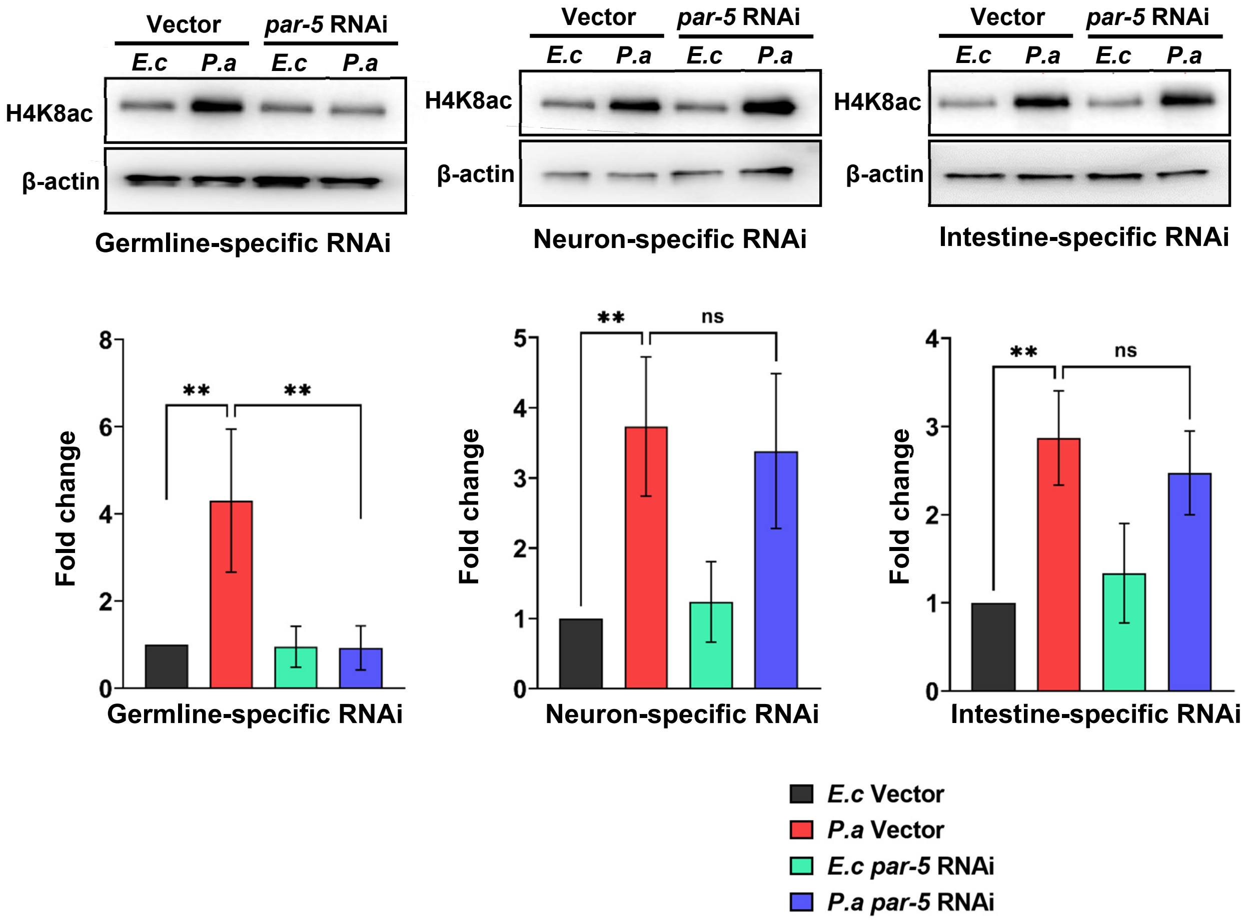

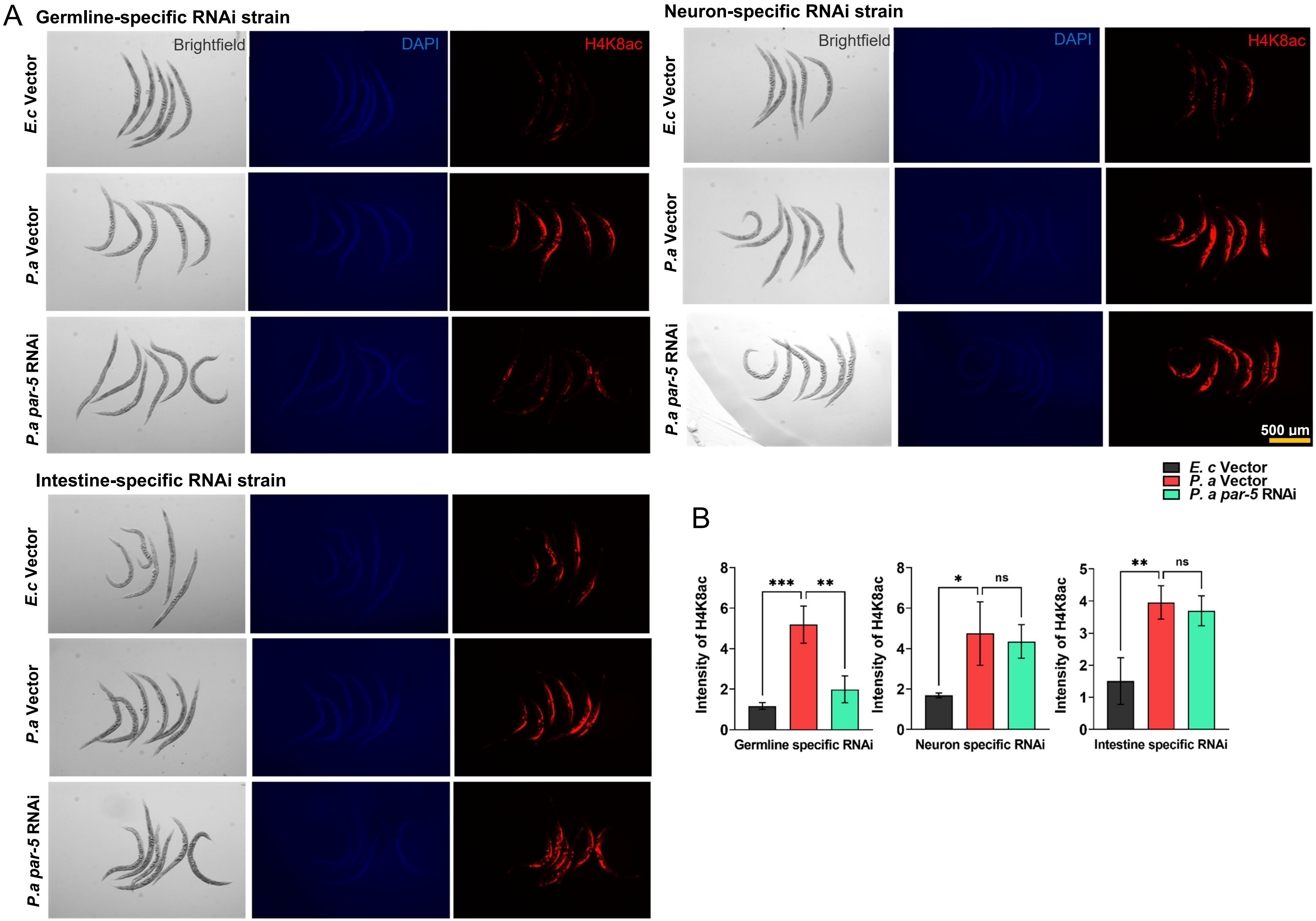

**S7 Fig.**

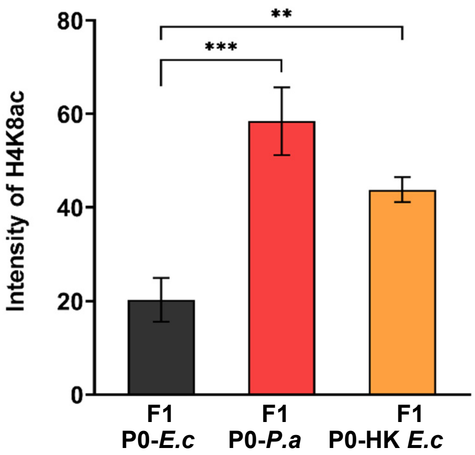
